# Functional mapping of the mouse *hairless* gene promoter region

**DOI:** 10.1101/244426

**Authors:** Eric G. Folco, Stefan Nonchev

## Abstract

The mouse *hairless* gene (*Hr*) encodes a protein of 127 kDa, acting as corepressor of nuclear hormone receptors. The Hairless protein (HR) is involved in the control of the cellular transition to the first hair cycle in adult Mammals. In its absence hair follicles disintegrate leading to a complete and irreversible hair loss with formation of cutaneous cysts. The hairless phenotype is therefore linked to defective proliferation and migration of the hair follicle stem cells apparently unable to respond to various signalling molecules. The *Hr* gene is expressed at high levels in skin and brain, and *hairless* transcripts were detected in gonads, thymus and colon. Although the patterns of *Hr* expression appear to be spatially and temporally regulated, very little is known about the molecular basis of the transcriptional control underlying *Hr* gene function. In this work we determine the precise transcriptional initiation start site of the mouse *Hr* gene and identify a new 1,1 kb cis-control element (RE1) that encompasses the promoter region and is able to drive luciferase reporter expression in skin and brain derived cell lines. We performed a deletion analysis and explored functionally regulatory motifs within this fragment to show that the role of this upstream regulatory region is linked to the presence of TRE and VDRE binding sites. We find that a TRE situated at –300 bp from the cap site is essential for gene expression in both skin NIH 3T3 and GHFT1 cells, while a VDRE positioned 94 bp upstream of the TRE modulates reporter expression specifically in skin derived cell lines. In addition, we define a novel cis-regulatory motif UE60, situated at the 5’-end of RE1 and likely to interact with both TRE and VDRE. Our data complete previous results on the possible existence of an autoregulatory pathway, implicated in *Hr* gene regulation. Taken together these findings reveal a complex molecular network that potentially links several signalling pathways in hair follicle formation. We discuss the organisation of the regulatory modules in the mouse *Hr* gene upstream DNA sequences in the light of the high homology of this region in mouse, rat and human.

## Introduction

In Mammals the hair follicle cycle is controlled by a complex network of genetic interactions where the hairless (*Hr*) gene plays a crucial role to maintain stem cell mediated hair growth and regression. The structure and expression of *Hr* was thoroughly analysed in mouse, rat and human [1], [2], [3], [4]. It was established that the Hairless protein (HR) is a nuclear receptor corepressor able to regulate gene transcription by its direct association to VDR, TR, ROR, as well as by its involvement in the control of Wnt signalling pathway and *Hox* gene expression [5], [6], [7], [8], [9], [10], [11] [12]. The *Hr* and *Vdr* genes are coexpressed in cells of the hair follicle, their mutations in mice and human share a common skin phenotype and their protein products interact physically in a way that this association drastically represses VDR mediated transactivation [8], [13], [14], [15]. The *Hr* gene is regulated by the thyroid hormone, the Hairless protein binds to TR *in vivo* and, at least in the brain, HR represses the transcription of TH-responsive genes by recruiting a particular complement of HDAC activities [16], [7]. Surprisingly, the majority of these elegant studies did not provide enough details on the *Hr* gene regulation itself, leaving open the question of a possible autoregulatory pathway that would function in various physiological contexts.

In the rat a TR/RXR binding site was mapped within a 106 bp fragment, 9 kb upstream of the first exon. A 23 bp sequence including an imperfect direct repeat of the type DR+4 was shown to act as a TRE [5]. A sequence matching a type DR+4 TRE was found at 2,6 kb upstream of the mRNA cap site in the human hairless gene and a 3 kb fragment containing this element was cloned with luciferase gene reporter and tested in absence and presence of T3 in brain and skin derived cell lines [17]. This study tackled the function of human *Hr* gene promoter fragments in specific cell types illustrating the complexity of differential transactivation mechanisms controlling gene expression in various tissue contexts. Indeed, the upstream sequences of the *Hr* gene include regulatory elements and putative transcription factor binding sites that were poorly described in rats and humans and never addressed in the mouse.

Here we report results on the fine structure of the mouse *Hr* gene promoter region. We mapped the precise *Hr* gene transcription start site and identified a one kb regulatory element (RE1) essential for promoter activity and harbouring consensus binding sites for TR and VDR. Epitope tagged VDRs and TRs were used to test the functionality of both TRE and VDRE by electrophoretic mobility shift assays (EMSA), and confirmed the ability of the putative sequences to bind specifically their transcription factors. Using a series of deletion derivatives, we performed functional analysis in skin and brain derived cell lines in order to narrow down the minimal regulatory motifs underlying context specific activity. We show that thyroid hormone and vitamin D are both able to boost RE1 driven reporter expression in the respective cellular context. Our data suggest that the consensus TRE, which is conserved in Mammals, but is a part of differently organised regulatory modules, is absolutely essential for *Hr* gene promoter activity. We identified a new 60 bp regulatory sequence (UE60) at the far 5’-end of the RE1, able to interact with the TRE and control a correct physiological response to hormone treatment. The role of the two VDREs identified appears to be much more subtle and suggest that these regulatory motifs contribute to the specific modulation of the promoter activity in skin derived fibroblasts. Taken together our data allow to identify new DNA control sequences in the promoter of the mouse *Hr* gene and shed more light on the molecular mechanisms of a putative autoregulatory pathway underlying its tissue specific transcription.

## Results

### Initiation start site of the hairless RNA transcription

Although the *Hr* gene structure and expression in mouse, rat and human have been thoroughly analysed in a number of laboratories, quite surprisingly, the gene transcription start site and the fine structure and properties of the immediate promoter region have not been clearly determined. Therefore we firstly set out to generate northern blots from various tissues and cell lines and hybridise them with a 1172 bp fragment probe generated by XbaI/HindIII digestion of mouse *Hr* cDNA cloned in pSK based vector. We detected a major transcript of 7,5 kb in skin, brain and heart RNA. No expression was found in the liver (Figure 1A). In addition to the organs, RNAs were extracted from a number of cultured cell lines. It is noteworthy that only the GHFT1 cells originating from the pituitary gland were expressing the *hairless* gene. Probing RNA from HaCat cells produced one weak and apparently non-specific band of about 5 kb, while no fragments were seen on the blots of the other cell lines (HaCat, HEK293, Cos, NIH 3T3 and HeLa) used (Figure 1A and not shown). As the length of the cDNA of the mouse *hairless* gene was confidently determined to be of about 4017 bp [1], we estimated that the precise site of initiating of RNA synthesis should be positioned at 4000 bp upstream of the ATG in the second exon of the mouse *hairless* gene. We then performed a primer extension analysis on total RNAs from several mouse organs at different developmental stages, as well as from various cultured cell lines in order to identify the *hairless* gene transcription start site. Using an end-labelled nucleotide of 25 bp (see Figure 2 and Material and Methods) designed to be complementary to a most 5’-end of the putative transcript, we produced in all the samples analysed a clear single stranded cDNA fragment of 645–660 nucleotides which allowed us to define more precisely the 5’-end of the transcript (Figure 1B). In order to elucidate ultimately the position and identity this promoter sequence, we designed another primer extension assay and obtained in skin and brain a band of 169 bp allowing to define precisely the sequence where the cap site is positioned. A manual sequencing reaction was then run along this second primer extension experiment in order to help determining the exact base pair and confirm the identity of the initiating guanine nucleotide (Figure, 1c). These combined mapping approaches allowed therefore to define for the first time the precise promoter domain of the mouse *hairless* gene and enabled us to start deciphering specific regulatory elements in the immediate upstream sequences in and around the promoter region (Figure 2).

**Figure 1.**
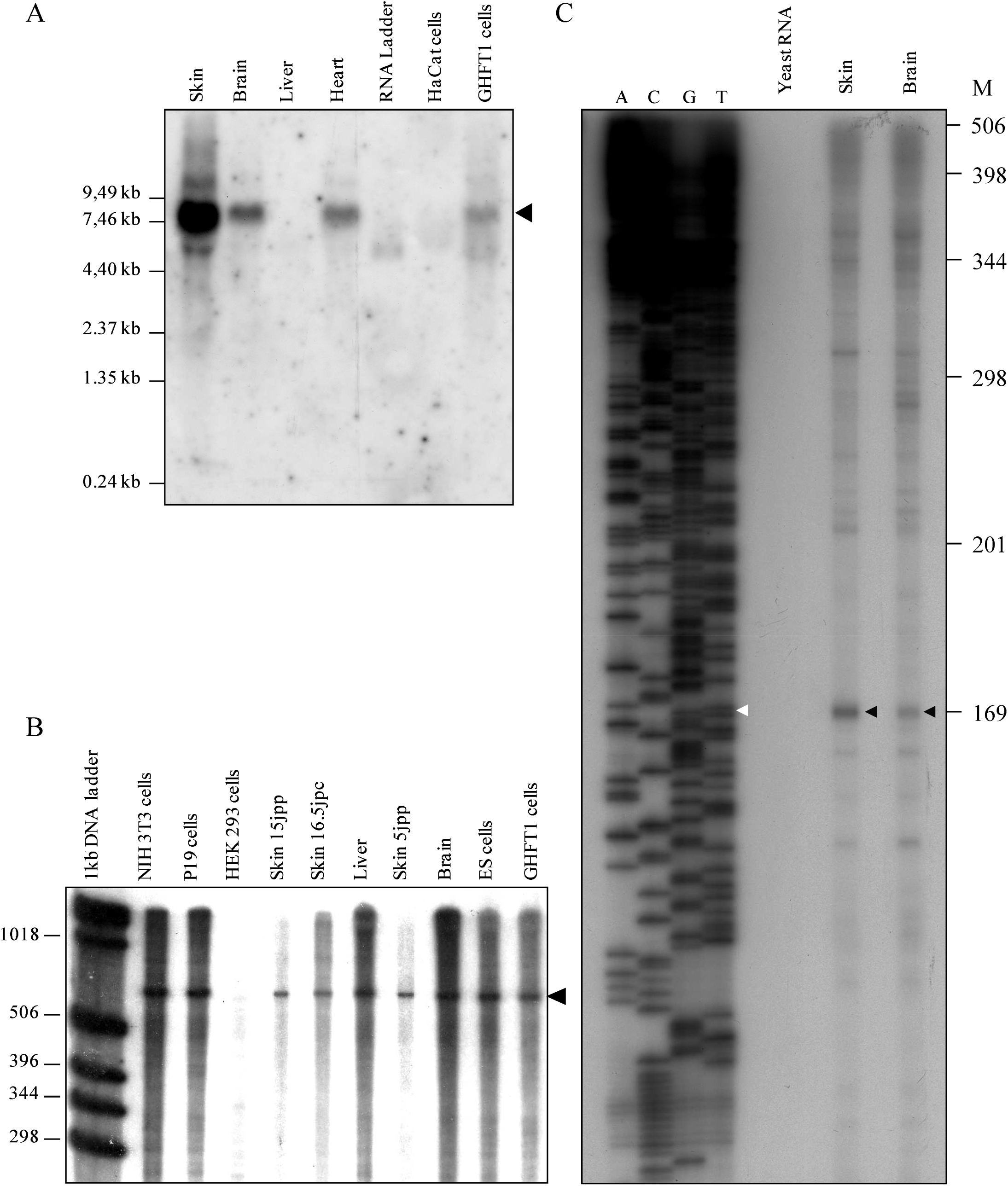
Determining the cap site of the mouse *hairless* gene. A. Expression of the Hr gene in various tissues and cultured cell lines. A specific transcript of 7,5 kb is detected by Northern blot in skin, brain and heart, as well as in pituitary gland derived GHFT1 cells. B. Primer extension analysis with a 5’ specific oligonucleotide. Note the 645–660 bp fragment produced in different organs and cell types. C. A second primer extension reaction yielded a 169 bp product, run along with a sequencing gel that allowed to identify precisely the cap site of the mouse *Hr* gene.

**Figure 2.**
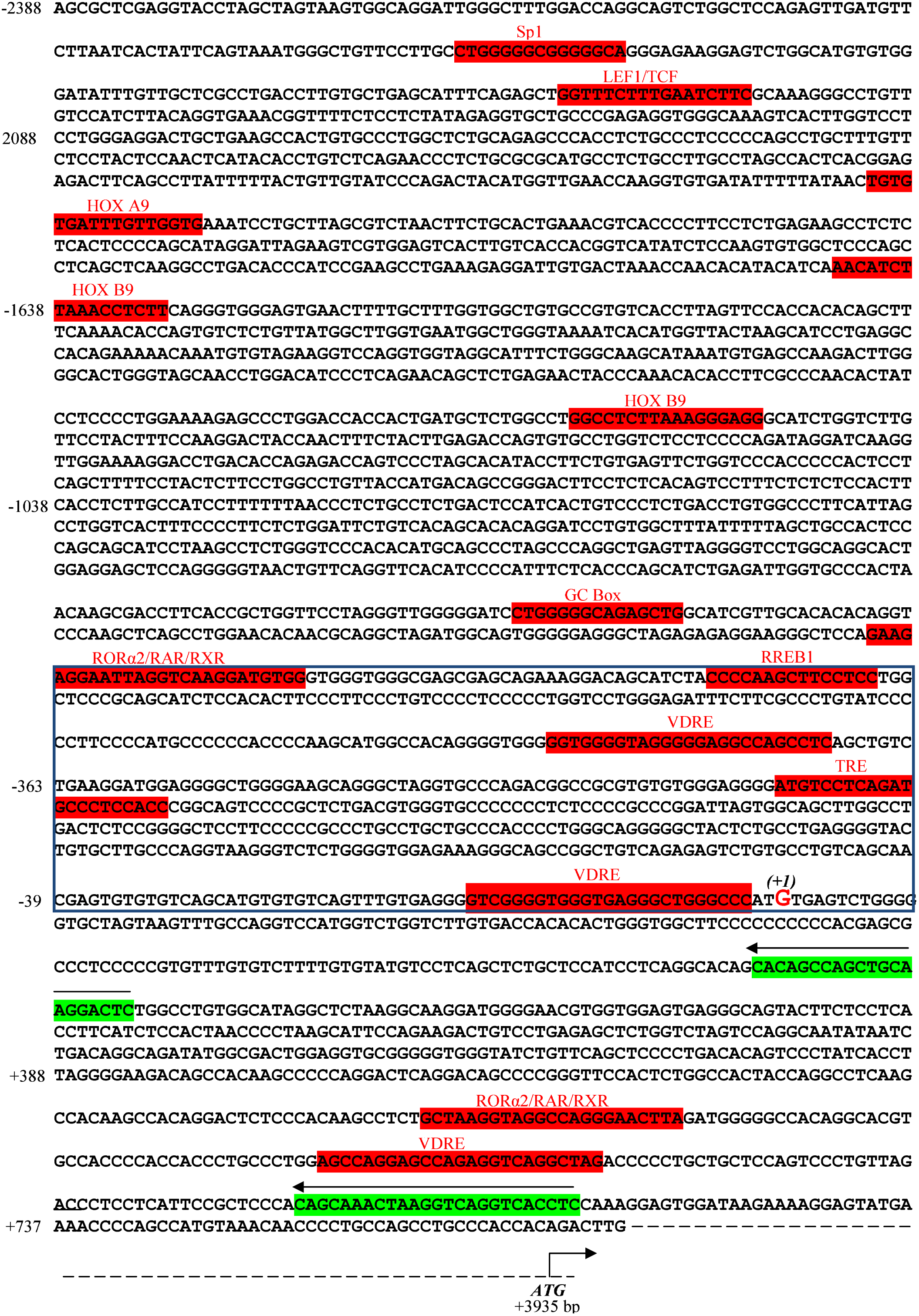
Nucleotide sequence (-2388 bp to + 737 bp) containing the mouse *Hr* gene promoter region. The cap site of the mouse *Hr* gene is situated 4 kb upstream of the HR protein translation start site (in the second exon of the mouse *Hr* gene). The oligonucleotides used for primer extension and sequencing reactions are shown in green. Putative transcription factor binding sites are coloured in red. The 1,1 kb regulatory element essential for the mouse *Hr* gene promoter function is dashed in blue. The punctuated line designates the 3200 bp separating this sequence from the mouse Hr gene ATG.

### Identification of DNA sequences essential for the Hr gene promoter activity in cultured cell lines

To test the regulatory potential of the region, we cloned a large 3 kb fragment encompassing the *Hr* gene promoter in a vector coupled to the luciferase reporter. Based on the fact that *Hr* is expressed strongly in skin and brain, we explored the ability of this big fragment to drive basal reporter expression in two different types of cell lines as models of transcription activity in fibroblasts (NIH 3T3) and pituitary gland (GHFT1). We then created a series of deletion derivatives and identified elements absolutely essential for basal transcription. As illustrated on Figure 3, construct 1 representing the whole 3 kb fragment (Figure 3A) contains elements able to promote high levels of reporter expression in both cell lines. Note that the levels of this activity are comparable in NIH 3T3 and GHFT1 cells (Figure 3B, 1). When this fragment was divided in two elements of roughly equal size we found out that construct 2 (nt –2388 to nt −769) was unable to drive reporter expression, while construct 3 (nt −837 to nt +461, Figure 3A) insured robust reporter activity of comparable levels in fibroblasts and neural cells (Figure 3B, 2 and 3). To monitor the efficiency of the border sequences of constructs 2 and 3, we generated constructs 4 (nt −2388 to nt −491) and 5 (nt –553 to nt +461). Here again, we established that this 1,1 kb long 3’ part of the whole fragment (construct 5) potentiated extremely high levels of luciferase expression especially as far as NIH 3T3 cells are concerned (Figure 3B, 5). In the same time, the whole 2,0 kb 5’-sequence (construct 4) was not efficient to govern reporter function neither in skin nor in brain derived cells (Figure 3B, 4). With constructs 6 (nt –837 to nt-491) and 7 (nt −1747 to nt –769, Figure 3A) we tested the role of short sequences adjacent to the 5’-end of the essential promoter element to find out that they were efficient in none of the cell lines (Figure 3B, 6 and 7). With construct 8 (nt – 2388 to nt −1575, Figure 3A) the effect of sequences remote to the promoter core was addressed and as it can be seen on Figure 3B, 8, this sequence is also unable to guarantee luciferase activity in the tested cell populations.

**Figure 3.**
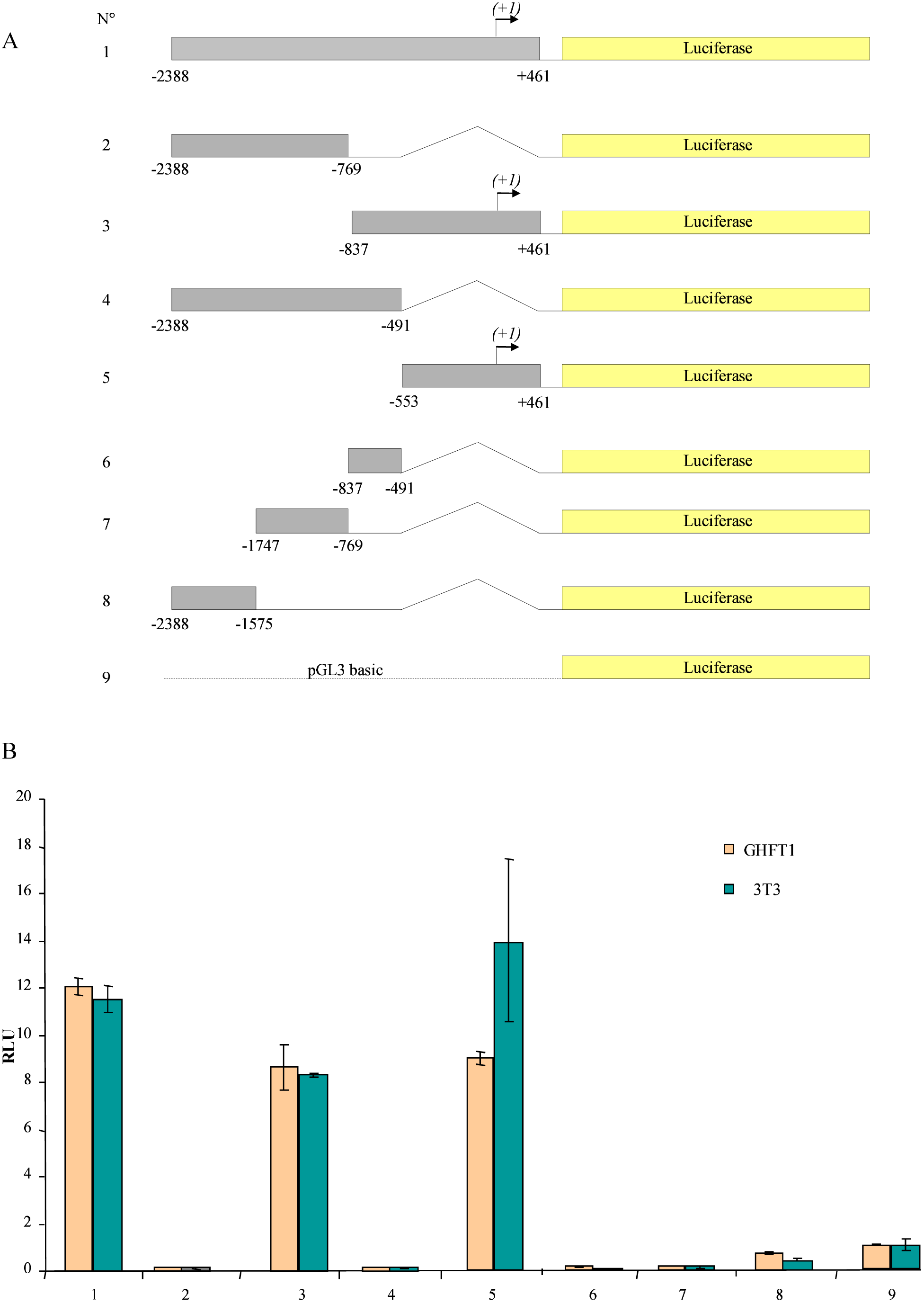
Identification of the RE1 (Regulatory Element 1) by deletion analysis. A. Deletion derivatives (constructs 1 to 8) of the 3 kb upstream region shown in Figure 2 were cloned in the pGL3 luciferase reporter vector (construct 9). B. Luciferase reporter expression in transfected NIH 3T3 (green) and GHFT1 (orange) cells. See Material and Methods for details of the transient transfection assay. Note that constructs 1, 3 and 5 are able to drive high levels of reporter activity, while the derivatives lacking the RE1 (constructs 2, 4, 6, 7 and 8) do not sustain luciferase transcription.

Taken together the data of this deletion analysis suggest that the regulatory elements responsible to govern the *Hr* gene basal promoter activity are organised within a 1kb fragment we named RE1 for regulatory element number 1, encompassing the transcription start site and its immediately adjacent upstream and downstream sequences. As shown on Figure 2 and as previously stressed, careful examination of this sequence allows to spot a number of potential upstream factors binding sites. Among those we got particularly interested in the strong concensus response elements – one for the transcription factor thyroid hormone receptor (TRE, nt −300 to nt −279) and two for the vitamin D receptor (VDRE1, nt −395 to nt −371 and VDRE2, nt −27 to nt −3, Figure 2). Despite the fact that direct protein – protein interaction of HR with TR and VDR was substantially described and thoroughly analysed [5], [8], very little is known about the possibility of a regulatory loop in which TR and/or VDR would regulate the *Hr* gene transcriptional activity in various cellular contexts [17].

### The TRE and VDRE within RE1 form specific complexes when incubated with extracts from TR and VDR transfected cells

Having delineated the 1,1 kb essential promoter region RE1, we pursued gel retardation experiments in order to test the capacity of the RE1 located VDRE and TRE to form complexes with VDR and TR from whole cell extracts. The electrophoretic mobility shift assays (EMSA) using double stranded radio-labelled nucleotides incubated with protein extracts from NIH 3T3 cells transfected with Flag tagged VDR and TRα2 are illustrated on figure 4A. As illustrated by Figure 4A, lane 1, the first radio-labelled sequence (nt −527 to nt – 409) did not display any migration shift. In fact this motif does not contain putative TRE or VDRE binding sites. The second probe (nt –527 to nt –321) contained the nucleotide sequence of the lane 1 plus the first putative VDRE sequence (nt –395 to nt –371). As shown on Figure 4A, lane 2, a substantial retardation is noted with this fragment. The third oligonucleotide (nt – 527 to nt –260) encompassing both sequences 1 and 2 contains an additional fragment of 61 bp, harbouring the putative TRE binding site. Therefore, specific complexes 1 and 2 leading to the observed gels shifts were formed with both TRE and VDRE fragments (Figure 4A, lanes 2 and 3). Treatment of the transfected NIH 3T3 cells with vitamin D3 (lanes 4 to 6) or with thyroid hormone T3 (lanes 7 to 9), did not affect complex formation and gel shift patterns after migration. The components of these complexes are now being analysed in super-shift experiments using a panel of antibodies to probe the specificity and the stability of the complexes formed.

**Figure 4.**
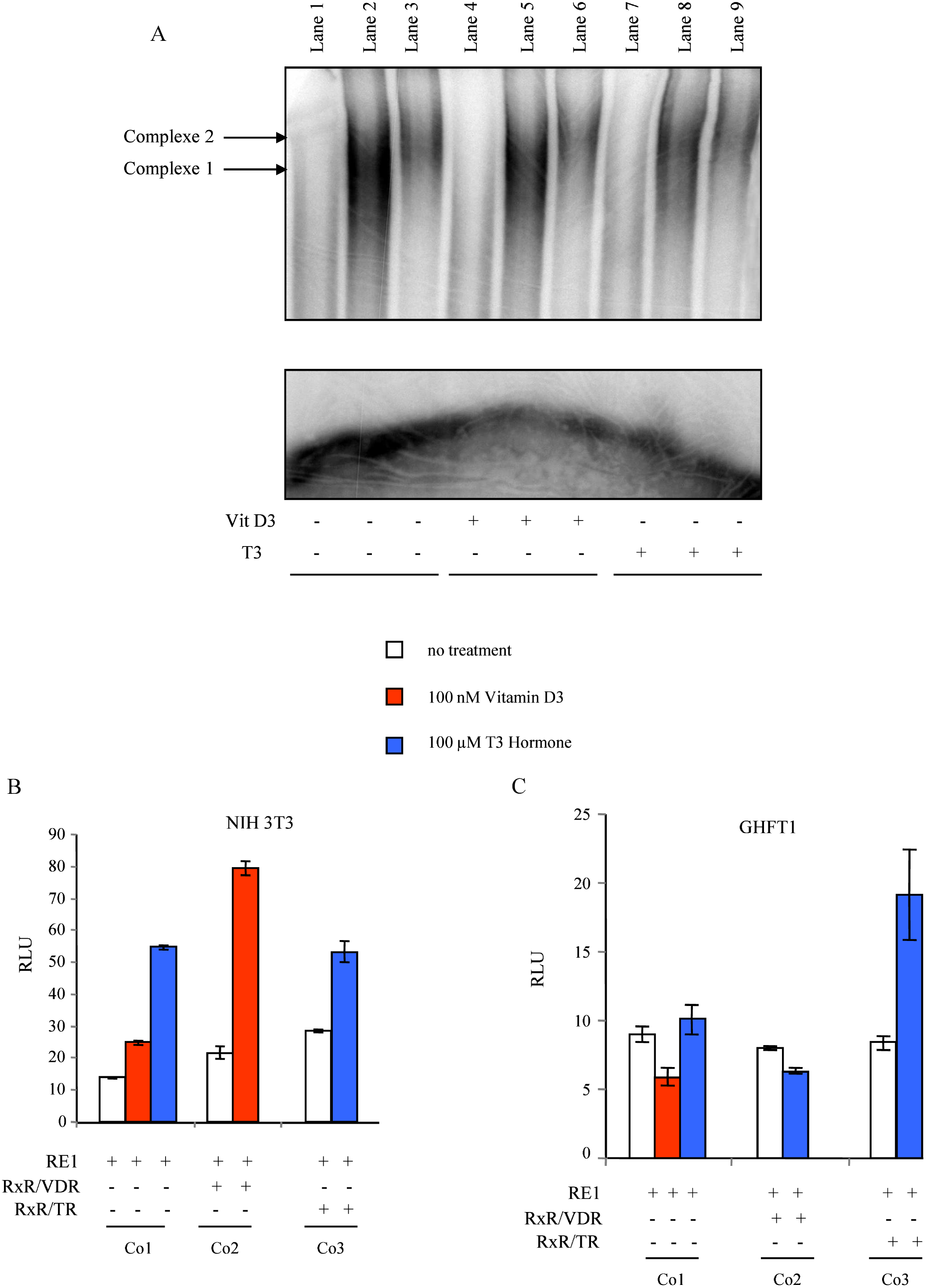
REI contains thyroid hormone and vitamin D response elements. A. An electrophorethic mobility shift assay (EMSA) with double-stranded radiolabelled VDRE and TRE specific nucleotides incubated with extracts from cells transfected with Flag-tagged VDR and TRα2. Notes the main complexes 1 and 2 arrowheads in lanes 2 and 3. Shifts are observed on lanes 2 and 3. No band shifts are detected in reactions with sequence 1. Transient cotransfection of NIH 3T3 (B) and GHFT1(C) cells with a construct of luciferase reporter coupled RE1 together with TR/RXR and VDR/RXR expressing plasmids in the presence or absence of the respective ligands, T3 thyroid hormone and vitamin D. Cotransfected cells were seeded in 60 mm plates at a density of about 3x10^5^ cells and then incubated overnight in the presence 100 µM T3 and 100 nM of vitamin D. The components of each transfection (Co1 to Co3) are indicated by + or −.

### Effects of Vitamin D and thyroid hormone on RE1 activity

To confirm the functional role of the newly identified regulatory motifs within the RE1 we designed cotransfection experiments in the cell lines that reflect promoter activity in skin and brain. NIH 3T3 and GHFT1 cells were then cotransfected with luciferase coupled RE1 together with either VDR/RXR or TR/RXR expression vectors. As expected, both vitamin D and the hormone T3 boosted basal reporter activity even in the absence of receptor coexpression in NIH 3T3 and in GHFT1 cell lines (Figure 4B and C). However, when VDR and TR were present in the transfected cells a robust increase of reporter expression was observed only when NIH 3T3 cells were treated with Vitamin D (Figure 4B, Compare Co2 and Co3). In contrast, when contransfection and ligand treatment were performed in GHFT1 cells, only the TR, but not the VDR, responded to ligand treatment by a massive increase of luciferase gene transcription (Figure 4C, Compare Co3 to transfections Co1 and Co2). Co-expression of VDR and RE1 had no effect on reporter activity that remained at similar levels with or without ligand treatment (Figure 4C-transfections Co1 and Co2). These data clearly suggest that despite the presence of both VDRE and TRE, the RE1 element could behave specifically in different cellular contexts. In particular, in brain derived cells it is sensible to thyroid hormone, but not to the vitamin D. This finding sheds more light on the molecular basis of this regulatory interaction and confirms previous observations that in developing brain, but not in the skin, the *Hr* gene is controlled by the TR.

### Deletion analysis of RE1 function

The next important question we addressed was the regulatory potential of the consensus VDRE and TRE and their interaction with other elements within RE1. We wanted to define even better the minimal motifs mediating specific ligands responses in NIH 3T3 and GHFT1 cell lines. To test this we generated a series of luciferase reporter coupled deletion derivatives from RE1 and explored their activity separately in NIH 3T3 cells and GHFT1 cells with and without respective ligand treatment. Surprisingly, in NIH 3T3 as well as in GHFT1 cells, deletion of the first 60 nt (Construct A, nt –504 to nt +461) reduced substantially the levels of reporter expression (Figure 5A, construct A and Figure 5B and C). Although the overall transcription activity remained robust (compared to luciferase reporter alone – the pGL3basic plasmid, construct G) in both lines, this sharp reduction in luciferase expression was in favour of an even more complex regulatory mechanisms governing the RE1 function and involving its most upstream nucleotides. We called this 5’ element UE60 (for upstream element of 60 nucleotides), examined the sequence for candidate transcription factor binding sites and pursued the exploration of its functionality in cotransfection experiments (see Figure 6A and B). Narrowing down the construct A by about one hundred bp, we obtained construct B −nt−423 to nt +461. This element encompasses the TRE as well as the two VDRE sequences (Figure 5A). The transient transfection of construct B in NIH 3T3 and GHFT1 cells did not modify luciferase transcription levels in the presence or absence of vitamin D and T3 respectively (Figure 5 b and c, construct B). A major observation in this analysis was the fact that deletion of VDRE1 (construct C, nt –368 to nt +461) did not affect at all the regulatory function of RE1 neither in NIH 3T3, nor in GHFT1 cells (Figure 5B and C construct C). In contrast, when the TRE was removed (construct D), a sharp loss of luciferase activity was recorded in both cell types (Figure 5B and C, construct D). The TRE seems therefore to be an essential part of RE1 and its function is not influenced by the cellular environment. In the same time the ligand treatment data suggest that thyroid hormone is efficiently boosting reporter expression of all constructs containing the TRE motif and does not affect reporter activity when this sequence is removed (Figure 5B and C – compare A,B and C to D, E, and F). Finally, with constructs D, E and F we observed a highly reduced reporter expression comparable to the one driven by the basic control luciferase plasmid pGL3basic in both transfected cell lines (Figure 5B and C – D, E, F). This result indicates that VDRE2, which is situated very near to the cap site has not the potential to activate transcription in these experimental conditions. To complete and refine this analysis we designed experiments with the element RE1 and two selected key constructs (constructs A and D from Figure 5A), mediating sharp changes in transcriptional activity and used them in cotransfection studies with TR/RXR dimmers and thyroid hormone treatment. It was not surprising to see that in NIH 3T3 cells, cotransfection of the TR/RXR with elements containing the TRE sequences (constructs RE1 and A) accompanied by T3 administration leads to levels of luciferase expression that were higher than those obtained without ligand treatment Figure 6A, Co1 and Co2). It was interesting to observe in this cell type that TR cotransfected RE1 responded much more vigorously to T3 than its deletion derivative construct A (Figure 6A, Co1 and Co2). This particular result confirms the importance of the 60 bp upstream motif previously identified. In addition, removing the TRE (construct D) abolished this difference and the overall levels of expression fell to a minimum activity with and without thyroid hormone T3 (Figure 6A, Co3). Intriguingly, in GHFT1 cells the same type of cotransfetion and ligand treatment restored the loss of reporter activity observed with construct A, compared to RE1 (Figure 6B, Co1 and Co2). The effect of the T3 is there but both RE1 and construct A displays the same regulatory potential in the presence or absence of the thyroid receptor (Figure 6B, Co1 and Co2). Once the TRE-less version is cotransfected, transcriptional activity and differences between treated and untreated cells are completely erased (Figure 6B, Co3). This group of data clearly show that independently of the type of cell line used and ligand treatments applied in the transfection experiments, the TRE motif plays a key role in RE1 function and *Hr* gene regulation.

**Figure 5.**
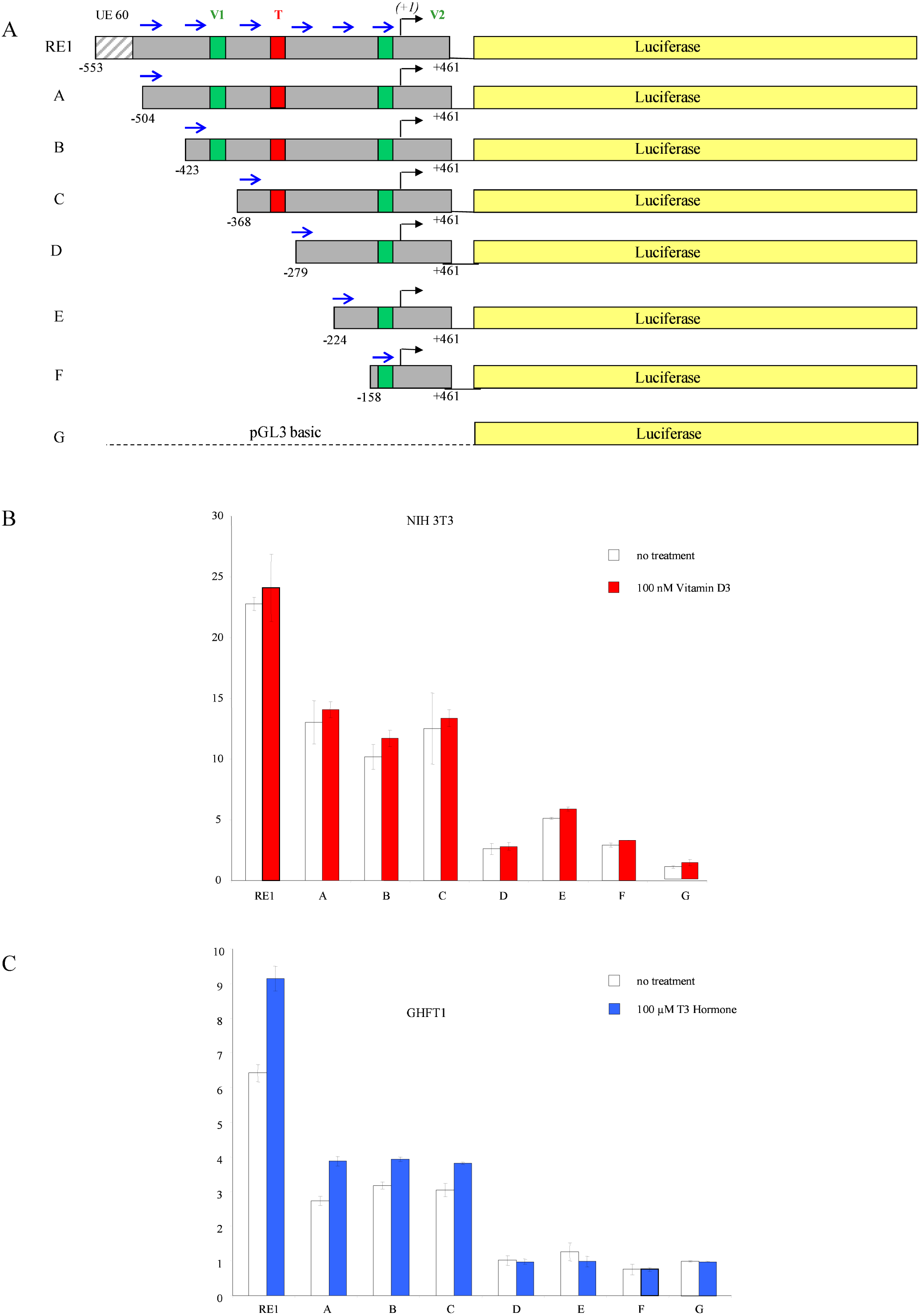
Functional analysis of the RE1 components. A. Serial truncations (A to F) of the 1,1 kb *H*r gene upstream regulatory element. The putative vitamin D response elements (VDREs) 1 and 2 are coloured in green (V1 and V2), while the presumptive thyroid hormone response element TRE (T) is shown in red o the respective constructs. The newly described regulatory element UE60 is marked in dashed. Construct G represents the pGL3 basic luciferase gene containing construct (see Material and Methods). B and C. Transient transfection assays of NIH 3T3 cells (B) and GHFT1 cells (C) showing the effect of RE1 regulatory motifs on gene transcription. Note the sharp loss of luciferase reporter activity when the UE60 element and the TRE are removed in the presence or absence of the respective ligand. Luciferase activity is expressed in Relative Luciferase Units (RLU).

**Figure 6.**
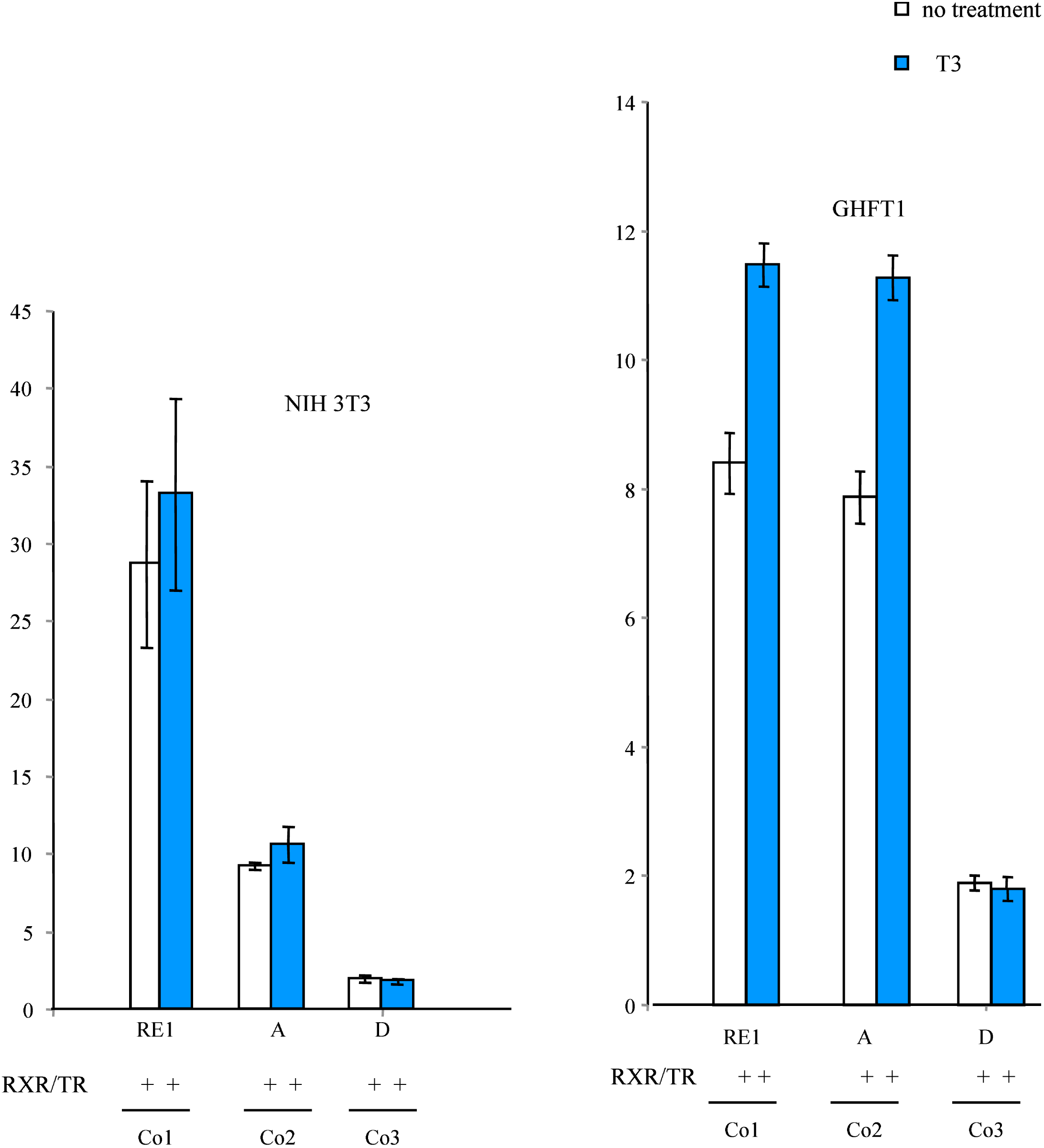
The TRE element is essential for RE1 function. A and E. Transient cotransfection assay in the presence or absence of thyroid hormone T3. Here NIH 3T3 cells (A) and GHFT1 cells (B) are contranfected with constructs RE1, A and D together with the TR/RXR receptors dimers. D. Reporter activity is dramatically reduced after removal of UE60 in cotransfection of NIH 3T3 cells, while it remains at high levels in the presenc or absence of T3 in cotransfected GHFT1 cells.

## Discussion

Although the mouse *Hr* gene is known for many years, its expression patterns have not been described until recently and its regulation remains unclear [18], [19] [20]. The upstream regions of this gene are largely unexplored and elements driving gene expression in a specific temporal and spatial manner were never identified. Indeed, prior experiments and analyses have stressed the fact that in rat and human the *Hr* gene regulation relied on upstream sequences containing proximal and distal predicted and predictable cis regulatory elements [5]. Based on the fact that the *Hr* is a thyroid hormone-responsive gene and that its protein product is a nuclear hormone receptor co-repressor, a partial analysis of its upstream regions in rat and human revealed functional TREs residing in the putative promoter.[21], [7], [17]. These data suggested that *Hr* is a target of hormone receptors signalling pathways and might be a part of an autoregulatory loop involved in the specific controls of downstream hormone responsive genes. In the present work we analysed in more detail the promoter region of the mouse *H*r gene. Taking into account the fact that many of the known cis-regulatory modules are made of clusters of phylogenetically conserved and repeated transcription factors (TF) binding sites, we started by restricting our analysis to modules located immediately and or moderately proximal to the *Hr* gene cap site. Using Northern blots, primer extension experiments and manual sequencing, we were able to define precisely the gene transcription cap site in the mouse at 3935 bp upstream of the ATG. Although the position of the *Hr* gene cap site in human and rat was previously inferred from human and rat library screenings with subsequent sequencing [5], [4], [17], here we defined for the first time the precise cap site, which allows to ultimately anchor the first nucleotide of the mouse *Hr* gene transcript and undertake a fine mapping of its promoter activity. We then tested systematically the regulatory potential of upstream sequences by transfection of deletion derivatives linked to luciferase reporter in skin and brain derived cell lines. A one kb regulatory element (RE1) was identified harbouring a number of putative transcription factors binding sites (Figure 2). A TRE motif of the type DR+4 was found at 300 bp upstream of the cap site and two potential VDREs that were not mentioned nor analysed before were noted within the essential RE1 sequence (Figure 2 and 7). We found that the TRE is essential for promoter activity in both cells lines used. In brain derived cells, however, this TRE is much more efficient than in fibroblasts (Figure 6A and B). When it is abolished, the promoter activity is completely lost (Figure 5A, B and C). Even if the first VDRE motif identified was not absolutely essential for the RE1 activity, its presence potentiates significantly the activation of the promoter specifically in the NIH 3T3 fibroblasts (Figure 5A, B and C). The second putative VDRE affects neither the whole RE1 activity nor that of its deletion derivatives. In contrast, the presence and the integrity of a 60 bp sequence (a fragment we call UE60) at the far 5’ site of the RE1 is critical for the overall activity of the *Hr* gene promoter as well as for the levels of reporter expression in response to ligand treatments in both cell lines; The RE1 element responded strongly to both vitamin D and T3 treatments It is important to note that the response to vitamin D is much stronger in NIH 3T3 cells, while the T3 response is more efficient in GHFT1 cells (Figure 4B and C). When the sequences upstream of the TRE are deleted, treatment with T3 is able to restore promoter activity specifically in GHFT1 cells, but not in NIH 3T3 cells (Figure 6A and B). Thus, in addition to the fact that the TRE is essential for the RE1 function in brain derived cells it interacts with the UE60 to potentiate a correct and physiological response to hormone treatment. The TREs consist typically of direct repeats sequences separated by a number of base pairs that can interact with hormone bound TR. As mentioned before, the functional TRE identified in this study is of the type DR+4 (GGG**GATGTC**CTCA**GATGCC**CTC) and is situated at 300 bp from the cap site (Figures 2 and 7). It is important to note that in the human and in rat DR+4 type of TREs were identified that varied in their repeated motifs as well as in their location with respect to the transcription start site of the *Hr* gene in these species [5], [17] and Figure 7. Like the TR, the vitamin D receptor (VDR) is a ligand-regulated transcription factor that recognizes cognate vitamin D response elements (VDREs) formed by direct or everted repeats of PuG(G/T)TCA motifs separated by 3 or 6 bp (DR3 or ER6)[22]. More than 900 vitamin D regulated genes are known in the mammalian genome and VDR binding was confirmed to several elements in vivo. VDREs lying within −10 to +5kb of the 5-ends were assigned to 65% of these genes. It was established that the effect of a nucleotide substitution on VDR binding in vitro does not predict VDRE function in vivo, because substitutions that disrupted binding in vitro were found in several functional elements [22]. In our hands the comparison of the two VDREs in the RE1 element shows that the VDRE1 has a fairly good core of consensus sequence TGAAGGA, while VDRE2, situated very close to the cap site displays less homology to the functional VDR recognition sites in various promoters [23].

**Figure 7.**
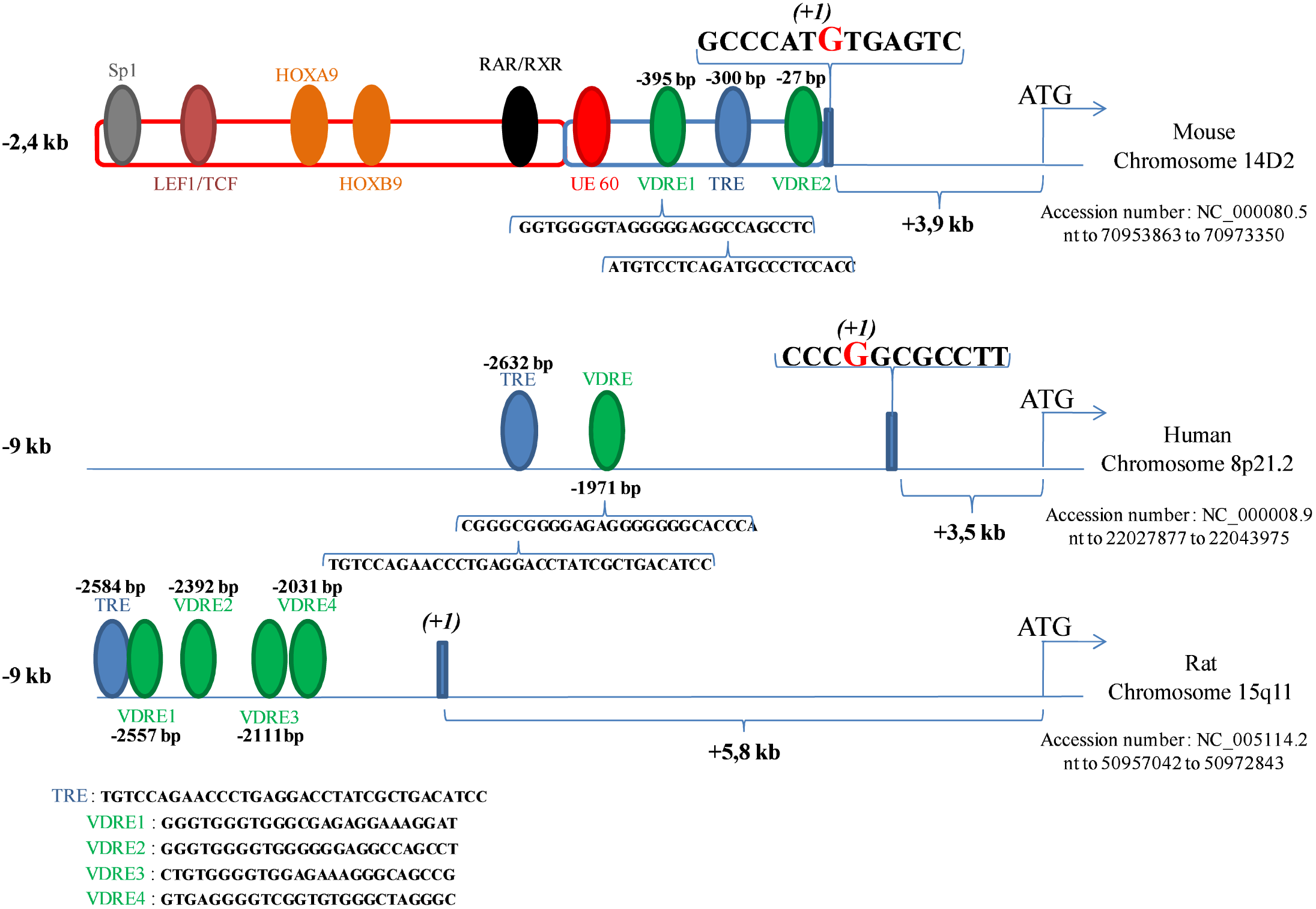
The *Hr* gene upstream regulatory modules are highly conserved in Mammals. The positions of a number of putative binding sites for transcription factors are indicated with respect to the *Hr* gene’s ATG in the three species. The precise locations of the promoters on mouse, rat and human chromosomes are marked. The binding sites identified and analysed in this work are shown in red (UE60), blue (TRE) and green (VDRE). The consensus sites for other upstream factors are presented in orange (Hoxa9 and Hoxb9), black (RAR:/RXR/ROR), purple (Lef1) and grey (Sp1).

The existence of an autoregulatory loop, by which activation of the *Hr* gene would contribute to the repression of downstream ligand responsive genes remains an open question in skin and hair follicle biology. The data of this work do not corroborate a previously postulated TRE mediated negative autoregulation that inhibits Hr promoter in keratinocytes but not in neuroblastoma cells [17]. By contrast, we found that thyroid hormone could enhance reporter expression from the RE1 in both skin and brain derived cells. Our results rather show that the T3 pathway is likely to exert its effect through of an yet unknown UE60 site binding upstream factor(s) and that VDREs modulate this interaction.

In fact, very little has been done to tackle the modular structure of the *Hr* gene regulatory sequences. In a pioneering study of Ahmad et al. 1999 a human genomic DNA sequence of 1058 bp 5’ of the *Hr* gene was analysed to establish the lack of TATA consensus sequences and to find by the Signal program putative binding sites for the transcription factors GATA-1, GATA-2, SP1 and AP1 [4]. Our data complete previous results on the thyroid hormone signalling responsive elements and shed more light on a possible interplay between TRE, VDREs and UE60 regulatory modules within and outside of the RE1 element. Our genomic blasts using the MatInspector program identified putative binding sites for Lef1/TCF and retinoic acid receptors known to mediate signalling that underlie fundamental processes of skin and brain development. Comparison of the upstream sequences suggests that these sites are again organised in a different way in mouse rat and human (Figures 7). Functional exploration testing the regulatory potential of the elements encompassing crucial upstream factors binding sites are now required and we started cloning the regulatory sequences identified by this study in LacZ and GFP reporter constructs in order to generate transgenic mice and decipher the temporal and spatial control of the *Hr* gene during development.

## Materials and Methods

### Primer Extension

For primer extension assays, a first antisense primer 5’-CAGGTGACCTGACCTTAGTTTGCTG-3’ complementary to the sequence from nt +626 to nt +651 (see figure 2, used in figure 1B), and a second antisense primer 5’-GAGTCCTTGCAGCTGGCTGTG-3’ complementary to the sequence from nt +148 to nt +169 (see figure 2, used in figure 1C) were labeled with [γ-^32^P]ATP and T4 polynucleotide kinase (NEB). Total RNA from mice tissues (skin, liver, brain, heart) and different cell lines (NIH 3T3, GHFT1, ES, P19, HEK 293, HaCaT) was prepared using TRIzol reagent protocol (Invitrogen) and RNAqueous kit (Ambion). 20 *µ*g of total RNA was hybridized with labeled oligonucleotide (10^5^ to 10^6^ cpm), qsp 50 *µ*L H_2_O. 0,1 volume of 3M sodium acetate and 2,5 volume of cold ethanol 100% were added for 1 hour at −20°C. The samples were precipitated with ethanol and resuspended in 20 *µ*L of a buffer containing 40 mM PIPES (pH 6.4), 1 mM EDTA (pH 8.0), 400 mM NaCl and 80% formamide. The samples were then denatured at 85°C for 10 min and let overnight à 42°C. The samples were precipitated with ethanol and resuspended in 50 mM Tris–HCl (pH 7.6), 60 mM KCl, 10 mM MgCl2, 1mM of each dNTP, 1 mM dithiothreitol, 20 units RNasin and 15 units SuperScript II reverse transcriptase (Invitrogen) and then incubated for 2 h at 42°C. The reaction was inactivated at 70°C for 10 min. The reaction mixture was treated with RNaseA, phenol and chloroform extracted, ethanol-precipitated, and resuspended in TE buffer (10 mM Tris (pH 7,5), 1 mM EDTA, 0,1 M NaCl). The products were analyzed by resolution on an 6% polyacryamide/8M urea sequencing gel and overnight autoradiography on Hyperfilm (GE Healthcare). The sizes of the resulting labeled primer extended products were inferred from their co-migration with a manual sequencing reaction, which was obtained using the same primer with a promoter-containing clone.

### Northern Blot

Tissues and cells were harvested and processed for isolation of total RNA as described above. Total RNA separated on formaldehyde agarose gels (15 µg/lane) and transferred onto nitrocellulose membrane by capillarity migration overnight. Pre-hybridization was carried out at 42°C for 2 h in pre-hybridization buffer (6X SSC, 50% formamide, 5X Denhardt’s, 1% SDS, and 200 µg/ml denatured salmon sperm DNA). Linearized cDNA probes (1172 pb obtained from pSK-Myc HR plasmid vector previously described (Brancaz-Bouvier et al, 2007) and cut by *Xba I-Hind III*) were radio-labeled by random-priming and heat denatured. Hybridization was carried out for 16 h at 42°C in pre-hybridization buffer. Blots were washed twice for 5 min at RT in 7X SSPE, 0.1% SDS, 1 time at 37°C in 1X SSPE, 0.5% SDS for 15 min and 1 time for 1 hour in 0.1X SSPE, 1% SDS at 65°C. Blots were exposed to Hyperfilm (GE Healthcare) at –80°C overnight.

### Cell culture and transient transfection assay

Cell lines were purchased from the American Type Culture Collection (ATCC) (Rockville, MD). Media and reagents for cell culture were purchased from Gibco (Invitrogen) unless otherwise indicated. Mouse fibroblast cell line (NIH 3T3) and mouse pituitary gland cell line (GHFT1) were maintained in DMEM high glucose (4,5g/L) medium containing L-glutamine, 10% fetal bovine serum, 100 units/ml penicillin, and 100 *µ*g/ml streptomycin at 37 °C under 5% CO_2_. Cells were plated on 12-well plates, each well containing 3x 10^5^. Following a 24-h incubation, cells in each well were transfected with 1,5 *µ*g of total plasmid constructs DNA (including promoter-reporter constructs, pRLTK-Luc, pSK as empty vector, etc.) using the SuperFect Transfection Reagent according to the manufacturer’s instructions (QiaGen). The pRLTK-Luc plasmid was co-transfected to normalize the variation in transfection efficiency. pRLTK-Luc encodes the *Renilla* luciferase, and its activity can be distinguished from that of the firefly luciferase encoded in pGL3b in the Dual-Luciferase Assay System (Promega). After 12 h of transfection, the cells were washed with phosphate-buffered saline and then maintained in fresh medium including or not a ligand treatments (100nM Vitamin D3 and 100*µ*M Thyroid hormone T3), depending of the experiments, for 12 h more prior to luciferase assay. In each experiment, the pGL3b plasmid was also transfected in separate wells to compare the specific activity of promoter-reporter constructs with the basic activity of the promoter-less plasmid.

### Dual Luciferase Activity Assay

Activities of the firefly luciferase and *Renilla* luciferase in a single sample were measured sequentially using the Dual-Luciferase Reporter Assay (DLRTM) system (Promega) according to the manufacturer’s instructions. Briefly, cells were rinsed twice with phosphate-buffered saline and then lysed in 200 *µ*l of Passive Lysis Buffer at room temperature for 15 min. 20 *µ*l of cell lysate was quickly mixed with 100 *µ*l of Luciferase Assay Reagent II in a luminometer tube. The light emission for the firefly luciferase was recorded immediately for 2s after a 3s pre-measurement delay using a Lumat LB 9507 Luminometer (EG&G Berthold). Subsequently, 100 *µ*l of Stop & Glo reagent was added to the same tube to inactivate the firefly luciferase while activating the *Renilla* luciferase. The light output from the *Renilla* luciferase was integrated under the same conditions. Variation in transfection efficiency was normalized by dividing the measurement for the firefly luciferase activity by that for the *Renilla* luciferase activity. The luciferase activity of a construct plasmid was expressed as relative to that of pGL3b in Relative Luciferase Units (RLU).

### DNA Constructs

cDNAs encoding the wild-type human VDR (gift from Dr. Hindrch Grosmeyer) and TRα2 (gift from Dr. Olivier Chassande) were subcloned into pcDNA3-Flag expression vector. Mouse genomic PCR was used to generate fragments with the following primers:

Primer 1: 5’-AGCGCTCGAGGTACCTAGCTAGTAAGTGGCAG-3’:nt −2388 to nt −2357

Primer 2: 5’-CACCACGGTCATATCTCCAAGTGTGGC-3’:nt –1747 to nt −1721

Primer 3: 5’-TGGAACTAAGGTGACACGGCACAG-3’ nt −1598 to nt −1575

Primer 4: 5’-GAGTTAGGGGTCCTGGCAGGCACTG-3’nt −837 to nt −813

Primer 5: 5’-GAAATGGGGATGTGAACCTGAACAG-3’nt −793 to nt −769

Primer 6: 5’-CGAGCAGAAAGGACAGCATCTACCC-3’nt −553 to nt −529

Primer 7:5’-GAAGTGTGGAGATGCTGCGGGAGCC-3’nt –515 to nt −485

Primer 8: 5’-CTGTGGCTTGAGGCCTGGTAGTGGCC-3’ nt +443 to nt +461

Construct 6: Primers 4 and 7 were subcloned in pGEM-T Easy (Promega), digested by *HindIII-SacI* and cloned in pGL3b (Promega). Construct 5: Primers 6 and 8 subcloned in pGEM-T Easy (Promega), digested by *HindIII-EcoRI* and subcloned in pSK II(-), digested by *KpnI-SmaI* and cloned in pGL3b (Promega). Construct 3: Construct 6 in pGEM-T Easy was digested by *HindIII-SalI* and subcloned in Contruct 5 in pSK. This new construct was digested in *KpnI-SmaI* and cloned in pGL3b. Construct 8: Primers 1 and 3 subcloned in pCR2.1 (Invitrogen), digested by *KpnI-EcoRV* and cloned in pGL3b (itself digested by *KpnI-SmaI*). Construct 7: Primers 2 and 5 were subcloned in pCR2.1 (Invitrogen), digested by *KpnI-EcoRV* and cloned in pGL3b (itself digested by *KpnI-SmaI*). Construct 2: Construct 8 was digested by *HindIII-StuI* and subcloned in Construct 7 in pCR2.1. This new construct was digested by *KpnI-EcoRV* and cloned in pGL3b (itself digested by *KpnI-SmaI*). Construct 4: Construct 8 was digested by *HindIII-StuI* and subcloned in Construct 7 in pCR2.1. This new construct was digested by *KpnI-SacI* and cloned in Construct 6 in pGL3b. Construct 1: Construct 3 in pGL3b was digested by *HindIII* and cloned in Construct 2 in pGL3b dephosphorylated. Construct 9 is pGL3b.

Deletion analysis of the RE1 fragment was performed by generating constructs A to F through PCR cloning of construct 5 truncations in pGL3b. The following primers were used:

Primer A: 5’-CGGGGTACCATCTCCACACTTCCCTTCC-3’:nt −504 to nt −485

Primer B:5’-CGGGGTACCACCCCAAGCATGGCCA-3’nt –423 to nt −406

Primer C:5’-CGGGGTACCTGTCTGAAGGATGGAGGG-3’nt –368 to nt −350

Primer D:5’-CGGGGTACCGGCAGTCCCCGCTCT-3’nt-279 to nt −263

Primer E:5’-CGGGGTACCAGCTTGGCCTGACTCTCC-3’nt –224 to nt −206

Primer F:5’CGGGGTACCTACTCTGCCTGAGGGGTA-3’nt −158 to nt −140

Construct A: Primers A and 8 were used in PCR (with construct 5 in pGL3b), digested by *KpnI-SmaI* and cloned in pGL3b. Construct B: Primers B and 8 were used in PCR (with construct 5 in pGL3b), digested by *KpnI-SmaI* and cloned in pGL3b. Construct C: Primers C and 8 were used in PCR (with construct 5 in pGL3b), digested by *KpnI-SmaI* and cloned in pGL3b. Construct D: Primers D and 8 were used in PCR (with construct 5 in pGL3b), digested by *KpnI-SmaI* and cloned in pGL3b. Construct E Primers E and 8 were used in PCR (with construct 5 in pGL3b), digested by *KpnI-SmaI* and cloned in pGL3b. Construct F: Primers F and 8 were used in PCR (with construct 5 in pGL3b), digested by *KpnI-SmaI* and cloned in pGL3b.

### Bioinformatic analyses

Genomic DNA sequences localized upstream the mouse, rat and human *Hr* gene (GenBank Accession Nos. NC_000080, NC_000008 and NC_005114 respectively) were taken from the ElDorado genome annotation database (Genomatix Software, Release 11/2007) and GenBank database (NCBI). The sequences were analyzed for phylogenetically conserved consensus motifs for transcription factor binding sites (TFBS) using the program FrameWorker (Genomatix Software) and BLAST (NCBI). The mouse, human and rat promoter region analysis was performed using MatInspector program (Genomatix Software, MatInspector Release professional 7.4.8.2, July 2007), [24].

### Electrophoretic Mobility Shift Assay

Plasmids pcDN3-Flag VDR and pcDN3-Flag TRα2 was transfected for 24h in 60 mm dishes using SuperFect Transfection Reagent (QiaGen) according to the manufacturer’s instructions. Treatment wad applied 12h after the transfection. Cells was scrapped with 1 mL PBS 1X, centrifuged at 2 000 rpm for 5 min at 4°C, flash-freezed in liquid nitrogen. Proteins was extracted with 3 volume of Buffer C (20 mM HEPES (pH 7.9), 25% Glycerol, 0.42 M NaCL, 1.5 mM MgCl_2_, 0.2 mM EDTA) for 10 min in ice and tube was centrifuged at 12 500 rpm for 5 min at 4°C. At the end, anti-proteases was added: 0.5 mM dithiothreitol, 0.5 mM pefabloc, 1 *µ*g/mL Leupeptin, 1 *µ*g/mL Pepstatin. Construct 5 in pSK was digested respectively by *Hind III-MscI (sequence 1 nt –527 to nt –409)*, *HindIII-EagI (sequence 2 nt –527 to nt –321)* and *HindIII-BmgBI (sequence 3 nt –527 to nt 260)* and each fragment was labeled with [γ^32^P]ATP and T4 polynucleotide kinase (NEB). Double-stranded DNA fragments used as probes were obtained by annealing complementary custom synthesized single-stranded oligonucleotides. Approximately 150 fmol of DNA was added to 10 *µ*g of nuclear extract in a final volume of 25 *µ*l containing 0.2 *µ*g of poly(dI-dC), 20 mM Tris-HCl (pH 8.0), 50 mM NaCl, bovine serum albumin (50 mg/ml), 1% Nonidet P-40, 1 mM EDTA, 10% glycerol, and 1 mM dithiothreitol. Following incubation at room temperature for 20 min, DNA-protein complexes were resolved by electrophoresis at room temperature through an 8% polyacrylamide gel (acrylamide/bisacrylamide 80:1) with 90 mM Tris borate (pH 8.5), 2 mM EDTA buffer. The gels were dried and subjected to autoradiography with PhosphorImager. Competition experiments included the addition of unlabeled DNA fragments to the reaction mix.

## Acknowledgements

We thank Dr Olivier Chassande for the TRα2 expression constructs and Dr.Henry Gronmayer for supplying the VDR and RXR plasmid constructs. This work was supported by the “Emergence” grant of the Region Rhône – Alpes and by the French “Fondation de la Recherche Médicale” (SN). EF had a PhD fellowship of the French Ministry of National Education. We appreciate the helpful discussion and encouragements from Drs John Sundberg, Catherine Thompson and Angela Christiano. We are grateful to Brigitte Peyrusse for her kindness, punctuality, and skilful artwork.

